# A Mendelian randomization study of glycemic and anthropometric traits and Parkinson’s disease

**DOI:** 10.1101/2020.03.31.017566

**Authors:** Sandeep Grover, Ricarda Graf, Christine Klein, Norbert Brüggemann, Inke R. König, Fabiola Del Greco M, Manu Sharma

## Abstract

**Background:** Impaired glucose and obesity are frequently observed in patients with Parkinson’s disease (PD), although it is unclear whether the impairment precedes or results from the neurodegeneration.

**Objective:** We aimed to assess whether glycemic and anthropometric traits can influence the risk of PD in 33,674 cases and 449,056 healthy controls using the Mendelian randomization (MR) framework.

**Methods:** We investigated causality with a two-sample MR approach in the European population to compute effect estimates with summary statistics from available discovery meta-analyses of genome-wide association studies (GWAS) on glycemic and anthropometric traits.

**Results:** We considered a threshold of p-value=0.0038 as significant after accounting for multiple testing, and p-value<0.05 was considered to be a suggestive evidence for a potential association. We observed a protective effect of waist-hip ratio (WHR) on PD (Inverse variance-weighted (IVW): OR _IVW_=0.735; 95%CI= 0.622–0.868; p-value=0.0003; I^2^ index=22.0%; MR-Egger intercept p-value=0.1508; Cochran Q test p-value=0.0003). The association was further retained after the exclusion of overlapping UK biobank (UKB) samples between the WHR and PD datasets (OR_IVW_=0.791; 95%CI=0.659–0.950; p-value=0.012; I^2^ index=13.0%; MR-Egger intercept p-value=0.733; Cochran Q test p-value=0.035). The sensitivity analysis provided suggestive evidence of an increased risk of PD on fasting glucose (FG) (β _IVW_=0.0188; 95%CI=0.0062–0.0313, p-value=0.0055; I^2^ index=0.0%; MR-Egger intercept p-value=0.0957; Cochran Q test p-value=0.4555) and protective effect of PD on T2D (Weighted median effect: OR_WME_=0.946; 95%CI=0.9290.983; p-value=0.0051; Weighted mode effect: OR_MBE_=0.943; 95%CI=0.904–0.983; p-value=0.0116).

**Conclusions:** Our results showed that central or abdominal obesity may be protective against PD development, independent of glucose levels.

## Introduction

The lack of neuroprotective or disease-modifying therapy has considerably hampered the management of Parkinson’s disease (PD). However, several recent preclinical and clinical studies have shown the potential beneficial effects of type 2 diabetes (T2D) specific treatment in exerting neuroprotection against PD, possibly by modulating glucose homeostasis and body weight^1-5^. Traditionally, insulin has been implicated in the general hormonal regulation of glucose metabolism, as insulin crosses the blood-brain barrier to modulate brain energy homeostasis, with a minor contribution from internal neuronal secretion^6^. A recent study also reported significantly higher blood glucose in non-diabetic PD patients compared to healthy controls during an oral glucose tolerance test, with no significant increase in insulin levels^7^. The study further reported an association of higher blood glucose levels with a higher BMI. Recently, type 2 diabetes (T2D) – characterized by high blood sugar, insulin resistance, and low insulin sensitivities – was shown to be associated with higher motor scores in patients with PD^8^. Change in body weight is also long known to occur during the clinical course of PD and with its treatment. A handful of observational studies with highly heterogeneous epidemiological study designs have investigated the association of body weight with PD, showing conflicting results although weight loss appears to be a consistent finding in more advanced PD^9-12^. In summary, inconclusive evidence from observational studies suggest that several highly correlated glycemic and anthropometric traits could alter the risk associated with development of PD. This could be attributed to limited sample sizes, presence of inherent confounding and reporting bias in observational studies.

Genome-wide association (GWAS) or meta-analyses of GWAS often have larger sample sizes with adequate coverage of the human genome, making GWAS-based Mendelian randomization (MR) an attractive approach. MR has recently evolved as an alternative statistical approach that can, against potential confounding, judge potentially causal relationships between risk factors (e.g. altered glucose metabolism or body mass index) and an outcome (e.g. PD)^13^. In principle, MR allows the use of genetic variants as proxy representatives of exposure from one population to test an association with an outcome in a completely independent population. The approach mimics the randomization of exposure in randomized controlled trials (RCTs) and, thereby, addresses hidden confounding factors. To date, MR studies exploring the causal role of altered glucose or insulin homeostasis in PD are lacking. However, a previously published study explored the role of body mass index (BMI) on PD and showed a protective role of body mass index (BMI) (OR = 0.82, 95% CI = 0.69-0.98)^14, 15^. Most recently, the availability of GWAS datasets from the UK Biobank has further made it possible to take advantage of an increased power associated with a higher sample size by meta-analyzing it with previously existing large scale consortium datasets on various phenotypes of interest^16-18^.

In the present study, we expand the spectrum by assessing the impact and influence of several glycemic traits including 2-hour post-challenge glucose (2hrGlu), fasting glucose (FG), fasting insulin (FI), fasting insulin (FPI), homeostasis model assessment of β-cell function (HOMA-B), homeostasis model assessment of insulin resistance (HOMA-IR), glycated hemoglobin (HbA1c), Modified Stumvoll Insulin Sensitivity Index (ISI), and Type II diabetes (T2D) as well as anthropometric traits including body mass index (BMI), waist-hip ratio (WHR), waist circumference (WC), hip circumference (HC), adult height (AH) and birth weight (BW) on PD.

## Methods

### Study design and identification of datasets

We conducted a two-sample MR study using summary estimates to examine the lifelong effect of glycemic and anthropometric traits on the risk of PD in the European population. We reviewed the most recent meta-analyses of discovery GWAS datasets in the literature and identified genetic instruments that influence glycemic traits including 2hGlu, FG, FI, FPI, HOMA-B, HOMA-IR, HbA1c, ISI, T2D and anthropometric traits including BMI, WHR, WC, HC, AH, and BW ^16, 18-28^ (**Table 1)**. For the outcome dataset, we used the discovery cohort of a recent meta-analysis of GWAS on 33,674 PD cases and 449,056 controls^17^.

### Prioritization of genetic variants

We extracted significant SNPs from each GWAS dataset by employing a cutoff of 5×10^−8^. All SNPs with F-statistics <10 were further excluded for a possible violation of MR assumption I^13, 29^. A clumping window of 10,000 kb and linkage disequilibrium (LD; i.e. r^2^) cutoff of 0.001 was applied in the European population in the 1000Genome Phase 3v5 dataset to identify the leading SNP that represents each significantly associated locus^30^. If a specific leading SNP was not available in the PD dataset, a proxy SNP (r^2^ > 0.8) was identified by using the European population in the 1000Genome Phase 3v5 dataset, when possible. The statistical power to detect a causal association was estimated by the method described by Brion et al. ^31^. Specifically, we set the sample size of the outcome dataset to 482,750 with 7.498% as proportion of PD patients in the dataset, a continuous exposure with a variance ≥1% and a threshold p-value of 3.8 × 10^−3^ (see the section below).

### Effect estimation using MR and test of pleiotropy

We used the inverse variance-weighted (IVW) method with first order weights as primary method to investigate the direct causal role of glycemic traits and anthropometric traits on PD^16-28^. If a genetic instrument consisted of a single SNP, we used the Wald ratio along with the delta method to estimate the causal effect and standard error, respectively. We applied a conservative Bonferroni correction to account for the number of 13 independent tests (local significance level = 0.05/number of tests) for a global significance of 0.05. All other results from statistical tests are interpreted descriptively. We used the intercept deviation test with MR-Egger’s and MR-Egger intercept and the I^2^ index to evaluate the heterogeneity of Wald estimates. Additionally, we also estimated the Cochran Q statistic for the IVW method as well as Rucker’s Q’ statistics for the MR-Egger’s method to evaluated the heterogeneity^32-35^.

### Sensitivity analysis

We employed MR-Egger, weighted median (WME), and weighted mode methods (MBE) methods to check the reliability of estimates by relaxing some of the MR assumption, thereby allowing instruments with varying proportions of pleiotropic variants, as previously explained^33, 36-39^.

To avoid the overlapping of samples from UK Biobank, which has been included in recently published GWAS, we computed casual effect estimates by using PD datasets without UK biobank samples, as used in the previous study (9,581 PD cases and 33,245 controls)^35, 40, 41^.

For glycemic traits, a number of studies reported GWAS results from replication and/or pooled data (**Supplementary Table 1a**). We therefore compared our primary results with those using genetic instruments from these additional studies to explore the issue of the bias due to the winner’s curse, led by the selection of the instruments done from the same dataset (discovery meta-analysis) used for the causal effect estimation.

Given that BMI has been shown to influence the role of glycemic and anthropometric traits on several diseases, including PD^7^, we also estimated the effect of genetic instruments adjusted for BMI for 2hrGlu, FG, FI, ISI, T2D, and WHR to identify their overall influence on the causal effect estimates for PD. A summary of GWAS datasets used to study the influence of GWAS study design and BMI adjusted datasets is provided in **Supplementary Tables 1a and 1b** respectively.

We employed a leave-one and leave-one-group-out cross-validation approach to check the influence of outlier variants as well as that of variants known to be associated with confounders of the relationship between glycemic, and anthropometric traits and PD. We used the PhenoScanner database to identify potential pleiotropic genetic instruments that are known to be associated with potential confounders^42^. Finally, we conducted a reverse directional MR by identifying genetic variants representing proxy markers of PD using the same study.

## Results

### Prioritization of genetic instruments and power analysis

The depth of genomic coverage and number of individuals in different discovery GWAS datasets is provided in **Table 1**. The table further shows the variance explained by genetic instruments for different exposure datasets and availability of genetic instruments in the PD dataset, estimated by the formula 2×β^2^×EAF×(1-EAF), where β is the estimated genetic effect of the exposure and EAF is the corresponding allele frequency^43^.

Two glycemic exposures (HOMA-IR and ISI) were not further analyzed given the absence of significant variants; however, we analyzed them as outcome with the PD as exposure in the reverse causation investigation. Thus, the number of primary statistical tests was reduced to the investigation of the causal effect of seven glycemic traits and six anthropometric traits on PD, leading to 13 tests. The significance level was accordingly set to 0.05/13 = 3.8 × 10^−3^.

Our power analysis suggests that our study has ≈80% power to detect a true OR of 1.208 or 0.799 for PD per SD of the continuous phenotype assuming that the proportion of the continuous phenotype explained by the genetic instrument is ≥1% at a type 1 error rate of 3.8 × 10^−3^.

### Effect estimation and sensitivity analysis

The causal effect estimates of glycemic traits and anthropometric traits on PD are shown in **Table 2**, which also provides various measures to evaluate the robustness of the effect estimates. The summary data used to compute effect estimates and sensitivity analysis is further presented in **Supplementary Table 2**. Among all the glycemic and anthropometric traits, we found that a 1-standard deviation (SD) increase in waist-hip ratio (WHR) was associated with a 26.5% lower risk of PD in the European population (OR_IVW_=0.735; 95%CI=0.622–0.868 per 1-SD of WHR; p-value=0.0003; I^2^=25.9%; MR-Egger intercept p-value=0.1508; Cochran Q test p-value=0.0003) (**Table 2a**). We further observed a similar effect using the WME method (OR_WME_=0.810; 95%CI=0.721–0.911). The distribution of individual SNP-level effect estimates along with the effect estimates computed through different MR methods for the effect of WHR on PD are shown as scatter and funnel plots in **Figure 1**. After ruling out the effect of weak instrument bias on account of overlapping UKB samples, the protective effect of WHR on PD was retained suggesting the reliability of the observed findings (OR_IVW_=0.791; 95% CI=0.659– 0.950; p-value=0.012; I^2^=13.0%; MR-Egger intercept p-value=0.733; Cochran Q test p-value=0.035).

**Figure 1.**
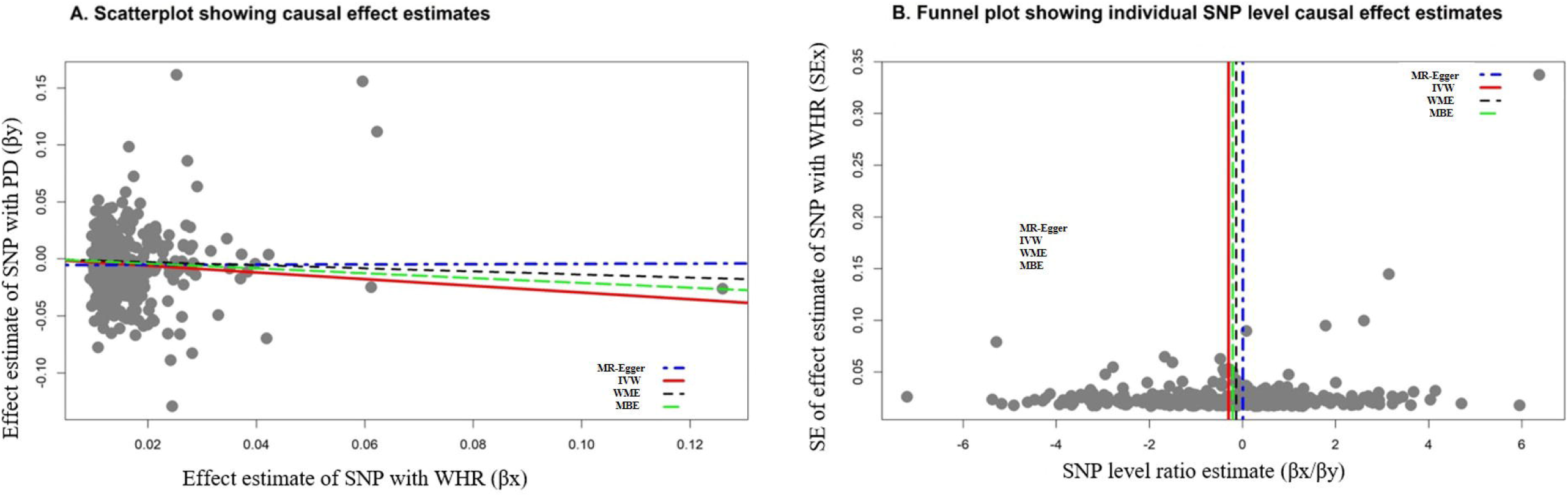
Graphical representation of causal association analysis and assessment of pleiotropy **A**. Scatterplot showing causal effect estimates computed using various MR methods for the association of Parkinson disease (PD) as exposure with Waist Hip Ratio (WHR) as an outcome. **B**. Funnel plot showing the extent of heterogeneity among the individual Wald ratio estimates for the association of Parkinson disease (PD) as exposure with Waist Hip Ratio (WHR) as an outcome.

Using the PhenoScanner database, out of 357 SNPs WHR associated SNPs employed in causal effect analysis; we identified a total of 127 pleiotropic SNPs that have been previously shown to be associated with non-anthropometric traits such as blood cell count, glycemic traits, lipid levels, and respiratory capacity (data not shown). In our sensitivity analysis that excluded these pleiotropic SNPs our instrument was weaker thus explaining the diminished protective effect estimate (OR_IVW_=0.801; 95% CI=0.640– 1.000; p-value=0.052). The leave-one out sensitivity analysis also failed to show influence of any single SNP, suggesting reliability of our findings (data not shown here). We further observed a loss of association when using genetic instruments for WHR which were adjusted for BMI, suggesting a potential role of BMI in influencing the causal association with PD.

The observed findings of the potential causal role of anthropometric traits was further confirmed by the absence of the causal effect of any of the glycemic traits, including FG and T2D on PD. This lack of association of glycemic traits further persisted when we used genetic instruments that were prioritized from a small proportion of moderately associated SNPs which were followed up in a pooled cohort for 2hGlu, FG, FI, and FPI (**Supplementary Table 3a**). Similarly, no association was observed for HOMA-B and HOMA-IR where genetic instruments were available for the replication cohorts. In addition, we did not observe any influence of the BMI-adjusted instruments that were available for 2hrGlu, FG, FI, and T2D, regardless of the GWAS study cohort that was used to extract the instrument (**Supplementary Table 3b**). With respect to other glycemic traits, we did not find genetic instruments for ISI. However, we were able to explore the causality using the single genetic instrument for BMI adjusted ISI phenotype which hinted at an association using genetic instruments prioritized from the discovery cohort only (OR_wald_ = 0.532, 95% CI=0.286-0.990; p-value=0.0464).

Lastly, we checked reliability of the observed relationships between various glycemic and anthropometric traits, and PD by conducting MR analyses in reverse direction, as shown in **Table 3**. We did not observe any causal effect of PD on WHR (β_IVW_=-0.0077; 95% CI=-0.0277–0.0122; p-value=0.4288; I^2^=84.1%; MR-Egger intercept p-value=0.8462; Cochran Q test p-value<0.0001). On the contrary, in our sensitivity analysis we observed the strongest effect on FG with a 1-log odds increase in genetic predisposition to PD being associated with 0.0188 mmol/l increase in FG concentration (β_IVW_ = 0.0188 per log-odds of PD; 95% CI=0.0062–0.0313; p-value=0.0055; I^2^=0.0%; MR-Egger intercept p-value=0.0957; Cochran Q test p-value=0.4555). Additionally, although IVW method failed to detect influence of PD on T2D (OR_IVW_=0.973; 95% CI=0.921–1.028; p-value=0.3258; I^2^=76.9%; MR-Egger intercept p-value=0.4711; Cochran Q test p-value=5.28×10^−11^), the genetic predisposition to PD was associated with approximately 5.0% lower risk of T2D using other methods (OR_WME_=0.946; 95% CI=0.930–0.973; p-value=0.0051; OR_MBE_=0.943; 95% CI=0.940–0.983; p-value=0.0116).

Our findings further motivated us to explore the triangulation relationship between the traits shown to be related to PD using the MR approach. Our data suggested a bidirectional causal relationship between T2D and FG as well as T2D and WHR (**Figure 2**) (data not shown). We further observed WHR as a potential risk factor for a higher FG with the absence of any effect of FG on WHR (**Figure 2**).

**Figure 2.**
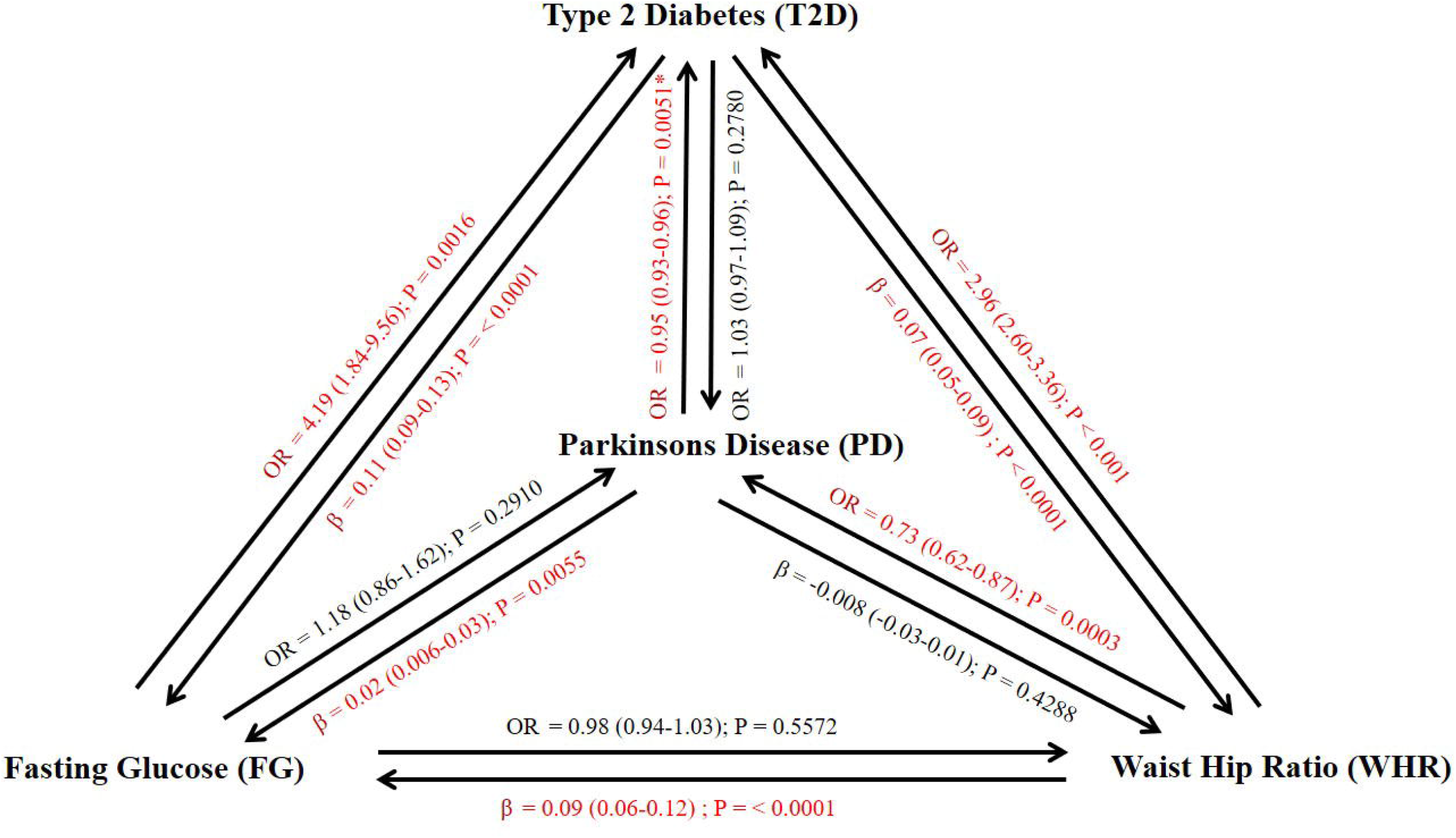
Graphical representation of causal relationship between Parkinson’s disease, glycemic and anthropometric traits. *Effect estimate computed using weighted median effect method (WME).

## Discussion

The present study using a two-sample MR approach aimed to understand the role of glycemic and anthropometric on PD and observed that an increase in WHR is a strong protective factor for PD. Furthermore, sensitivity analyses provided suggestive evidence, though not conclusive, of higher glucose tolerance and protection against T2D in PD patients.

Dopamine neurotransmission in the human brain is known to modulate the rewarding properties of food. Previous studies have further shown that dopamine receptors are under expressed in obese individuals, thereby initiating a feedback look to compensate for lower dopamine secretion^44^. It is, however, not known whether an altered dopaminergic metabolism in overweight individuals could influence the PD susceptibility. Several indicators have been used to measure overweight and obesity. While WHR and WC are predominantly used as measurements of central obesity, BMI is used as measurement of overall obesity. It has also been shown that WHR and WC may be regarded as better alternatives to BMI to measure obesity, especially in individuals with cardiovascular risk factors including T2D^45, 46^. Numerous observational studies have previously explored the association between both measures of obesity and PD with mixed results^9, 10^. A recent meta-analysis of ten cohort studies with 2706 PD cases showed an absence of association of BMI with PD^10^. In contrast, a recent nationwide health check-up data for the whole South Korean population comprising 44,205 incident cases identified risky association of abdominal obesity with PD (HR=1.13, 95% CI=1.10–1.16)^9^.

Using an MR approach, we observed a significant risk reduction of 26.5% with every one unit of SD increase in WHR, while not observing any role of BMI. Our results are in contrast with a previously reported protective causal association of BMI with PD, which observed a significant risk reduction of 18%^14^. The previous study however assessed 77 loci for exploring causal effect of BMI comprising 13,708 PD cases and 95,282 controls compared to 548 loci assessed in a pooled dataset of 33,764 PD cases 449,056 controls in the present study. The discrepancy in the number of prioritized loci between the two studies is attributed to the use of GIANT dataset on BMI in the previous study compared to pooled dataset of GIANT and UKB on BMI used in the present study. Another recent MR study prioritizing genetic instruments using the UKB dataset only and exploring the casual role of 401 exposures did not detect a casual role of BMI and WHR^15^. Nevertheless, they observed a consistent protective causal association of nine adiposity related traits with the strongest effect observed for arm fat percentage. In line with our findings, these studies collectively argue that the assessment of fat mass vs. fat-free mass e.g. by using body plethysmography reflects the underlying causative or protective factors more closely than body weight or BMI. Interestingly, a recent study reported a slight reduction of 1.12% in PD risk for every 1kg/m^2^ increase in BMI without any clear evidence of heterogeneity^47^. The study employed 23andMe dataset for both BMI and PD datasets and the absence of heterogeneity in the observed findings possibly suggest that a considerable heterogeneity observed in previous MR studies could be attributed to the clinical heterogeneity among the commonly employed IPDGC PD cases. Our study provided strong evidence regarding the role of WHR in PD, nonetheless, further studies are highly warranted to disentangle the protective role of WHR in PD.

The results observed in our present study provided further suggest a potential role of glucose metabolism in PD, and data obtained herein agree with the previously published epidemiological studies. For example, a recent study reported a significantly higher area under time curve (AUC) for the blood glucose levels in 50 non-diabetic PD patients compared to 50 healthy controls during a 75g oral glucose tolerance test (1187 ± 229 vs 1101 ± 201 mmol/min.; p=0.05), with no significant difference in AUC for blood insulin levels (6681 ± 3495 vs 7271 ± 6070 mmol/min.; p=0.57)^7^. The study also reported that higher blood glucose levels were associated with higher BMI (p-value<0.0001). Another recent longitudinal study identified high blood glucose as a risk marker for PD progression^48^. The 48-month follow-up study exploring the role of 44 clinical variables in 135 patients with early PD, identified high FG levels (p-value=0.013) and T2D (p-value=0.033), among several other factors as significant predictors of annual cognitive decline in PD. The study further observed significant differences in the baseline levels of glucose when compared to 109 healthy controls. Our results are henceforth in consent with these results suggesting that PD promotes dysregulation of glucose metabolism.

The relationship between PD and T2D as observed in our study is intriguing. We observed association only in sensitivity analyses. Thus, the results should be interpreted with caution. Nevertheless, several longitudinal studies have previously explored the influence of pre-existing T2D on the predisposition to PD with contradictory results. A prospective follow-up of 147,096 predominantly Caucasian participants in the Cancer Prevention Study II Nutrition Cohort from the United States found no association of the history of diabetes with PD risk (RR=0.88; 95% CI=0.62–1.25)^49^. Another study that comprised two large US cohorts – the Nurses’ Health Study (121,046 women) and the Health Professionals Follow-up Study (50,833 men) – observed similar results (RR=1.04;95% CI=0.74–1.46)^50^. In contrast, a follow-up study in 51,552 Finnish individuals demonstrated an increased incidence of PD among patients with T2D (HR=1.85; 95% CI=1.23–2.80)^51^. Most recently, a meta-analysis of four cohort studies (3284 PD cases and 32, 695 diabetes cases) confirmed the finding that the onset of diabetes was a risk factor for PD (RR=1.37; 95% CI=1.21–1.55)^52^. However, the same study reported the absence of an association in pooled populations of five case-control studies (6487 PD cases and 1387 diabetes cases; OR=0.75; 95% CI=0.50–1.11). In summary, findings of the association between T2D and PD have been highly heterogeneous and could be attributed to the varying age of onset of T2D and PD. Indeed previously published meta-analysis showed that the earlier onset of T2D before the onset of PD found to be a major risk factor for future PD patients^53^. Our close examination of GWAS datasets used in our study showed a remarkable variability in age of onset in PD and T2D subjects. For example, the majority of PD patients had an age of onset ranging from 48.9 to 71.2 years as compared to the average 52.5 years as an age of onset in T2D patients. Thus, it is conceivable that the earlier prodromal phase of PD, as compared to diabetes onset, could be protective against T2D in PD patients. Nevertheless, an in-depth clinical evaluation of PD subjects is highly warranted to further discern the role of T2D in PD etiology.

Despite our inability to stratify patients by the age of onset, our study has several strengths. We adopted a comprehensive approach that included several known markers of insulin metabolism. However, we observed that the genetic instruments for FI, HOMA-B, and HOMA-IR explained a very low amount of variance and, therefore, potential causation with PD might not be completely ruled out. An important limitation of this study could be the unavailability of individual-level data, which could have enabled us to confirm the absence of pleiotropic variants by using various potential confounding variables between WHR and PD. For instance, it is known that different markers of obesity show gender specific cut-offs and possibility of specific gender in influencing the observed causal estimates cannot be ruled out^54, 55^. Although we observed a suggestive reverse causal association of T2D with PD using different MR methods, we did not observe similar results with HbA1C, which is a known biomarker for prediabetes or diabetes. One of the reasons for this could be that the GWAS on HbA1c with 123,491 individuals from the general population was underpowered when compared to the GWAS on T2D that included 898,129 individuals^20, 21^. Another potential limitation could be the existence of overlapping UKB samples between WHR and PD datasets, which could have led to possible bias in our findings. Nevertheless, our sensitivity analysis demonstrates retention of the association even after the exclusion of UKB samples from the PD dataset, highlighting the robustness of our novel finding^56^. Lastly, we could not conduct a causal association analysis among different glycemic traits within PD patients.

Despite these limitations, our study represents one of the most comprehensive studies to date that has explored the potential causal role of glycemic and anthropometric traits on PD. Our analyses suggest that central obesity may play a role in conferring protection against PD. An extensive sensitivity analysis further suggested a possible role of PD in altered glucose metabolism independent of insulin activity. Furthermore, we showed that, despite high fasting glucose levels, PD patients may be protected against T2D. We further suggest the adoption of a cautionary approach when drawing clinical interpretations from the results of the current study, because additional lines of evidence may be generated, including the potential complex relationship of anthropometric, glycemic and PD with other unexplored traits.

## Author contributions

1. Research project: A. Conception, B. Organization, C. Execution; 2. Statistical Analysis: A. Design, B. Execution, C. Review and Critique; 3. Manuscript Preparation: A. Writing of the first draft, B. Review and Critique;

**S.G**.: 1A, 1B, 1C, 2A, 2B, 3A;

**R.G**.: 2A, 2B;

**F.D.G**.: 2C;

**N.B**.: 3B

**C.K**.: 3B

**I.R.K**.: 2C, 3B;

**M.S**.: 1A, 1B, 2C, 3B

## Acknowledgments

This study is, in-part, supported by the EU Joint Programme - Neurodegenerative Diseases Research (JPND) project under the aegis of JPND (www.jpnd.eu) through Germany, BMBF, funding code 01ED1406. Dr. Sharma is further funded by the Michael J Fox Foundation, USA Genetic Diversity in PD Program: GAP-India Grant ID: 17473. and also supported by the grants from the German Research Council (DFG/SH 599/6-1 to M.S.), MSA Coalition, and Michael J Fox Foundation. This study was also supported by grants from the German Research Foundation (Research Unit ProtectMove, FOR 2488) to I.R.K., F.D.G.,N.B.,C.K).

Data on PD GWAS was contributed by IPDGC team and downloaded from https://pdgenetics.org/. Data on glycemic traits were contributed by MAGIC investigators and downloaded from www.magicinvestigators.org. Data on T2D were contributed by DIAGRAM investigators and downloaded from www.diagram-consortium.org. Data on BMI, WHR, WC, HC and height were provided by GIANT consortium and downloaded from https://portals.broadinstitute.org/collaboration/giant/. Data on birth weight was provided by EEG consortium and downloaded from https://egg-consortium.org/.

## List of figures and tables

**Table 1**. Details of the discovery GWAS datasets that explored and prioritized genetic instruments used for direct and reverse casual analysis in the present study. The direct analysis was done using Parkinson’s disease (PD) as an outcome, and the reverse was done using glycemic traits and modifiable anthropometric traits as an outcome.

**Table 2**. Causal effect estimates using different Mendelian randomization (MR) methods and heterogeneity analysis of causal effect estimates for Parkinson’s disease (PD) by using various (a) glycemic traits as exposures. (b) anthropometric traits as exposures.

**Table 3**. Causal effect estimates using different Mendelian randomization methods and heterogeneity analysis of causal effect estimates for various (a) glycemic traits and (b) modifiable anthropometric traits by using Parkinson’s disease as an exposure.

**Supplementary Table 1a**. Details of follow-up genetic variants in the replication and pooled exposure GWAS datasets and prioritized genetic instruments used for the secondary analysis

**Supplementary Table 1b**. Details of exposure GWAS datasets adjusted for BMI and prioritized genetic instruments used for the sensitivity analysis

**Supplementary Table 2**. Harmonized summary effect estimates from exposure and outcome datasets used for the conduct of Mendelian randomization (MR)

**Supplementary Table 3**. Sensitivity analysis exploring the (a) influence of GWAS study design (b) BMI adjusted traits on causal effect estimates

## Financial disclosures

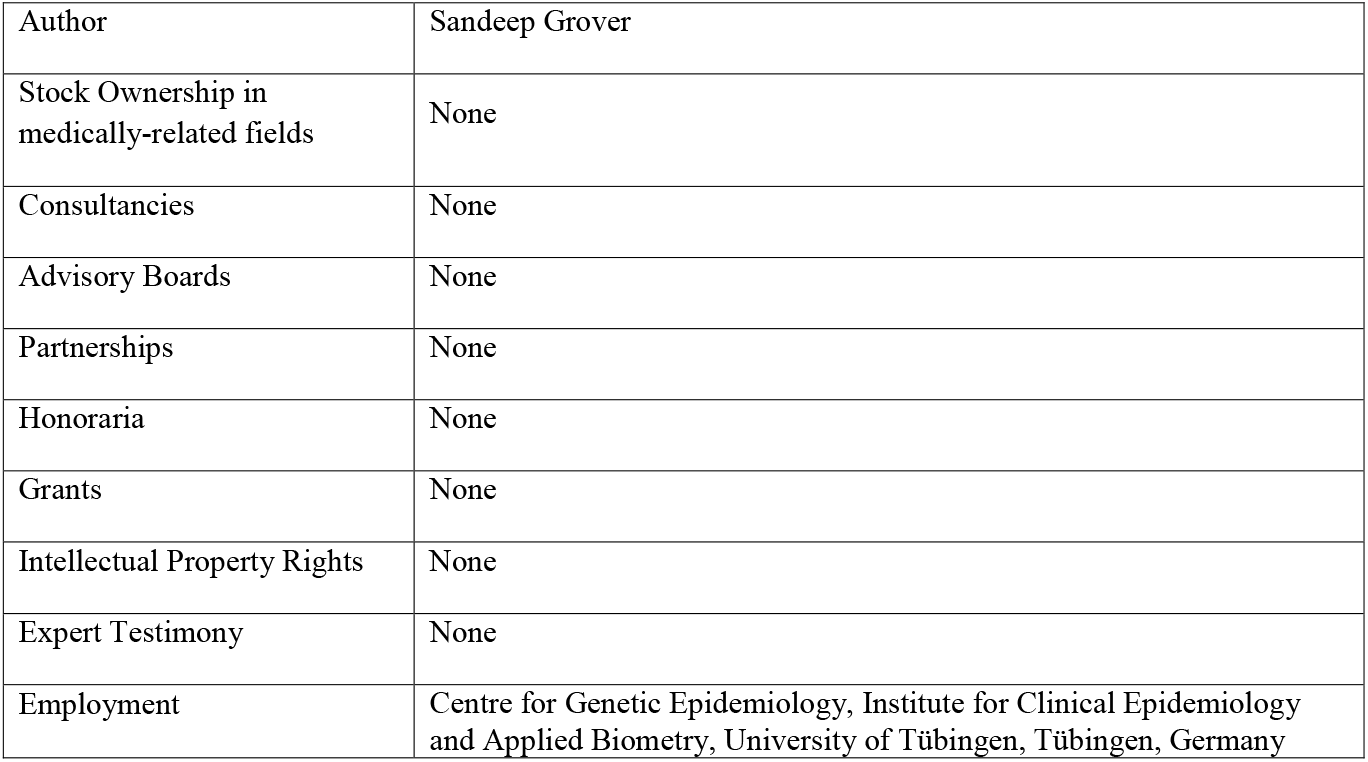

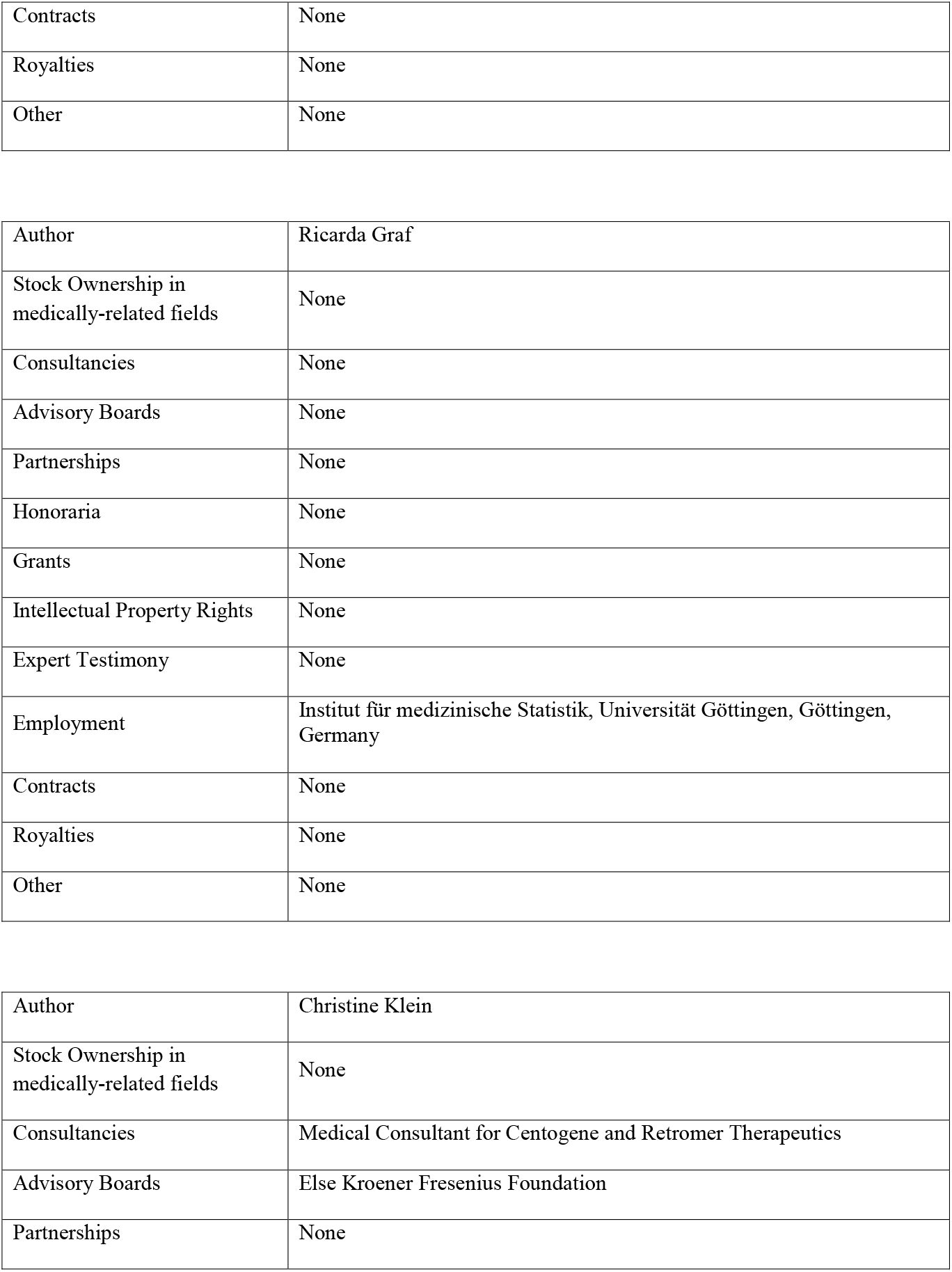

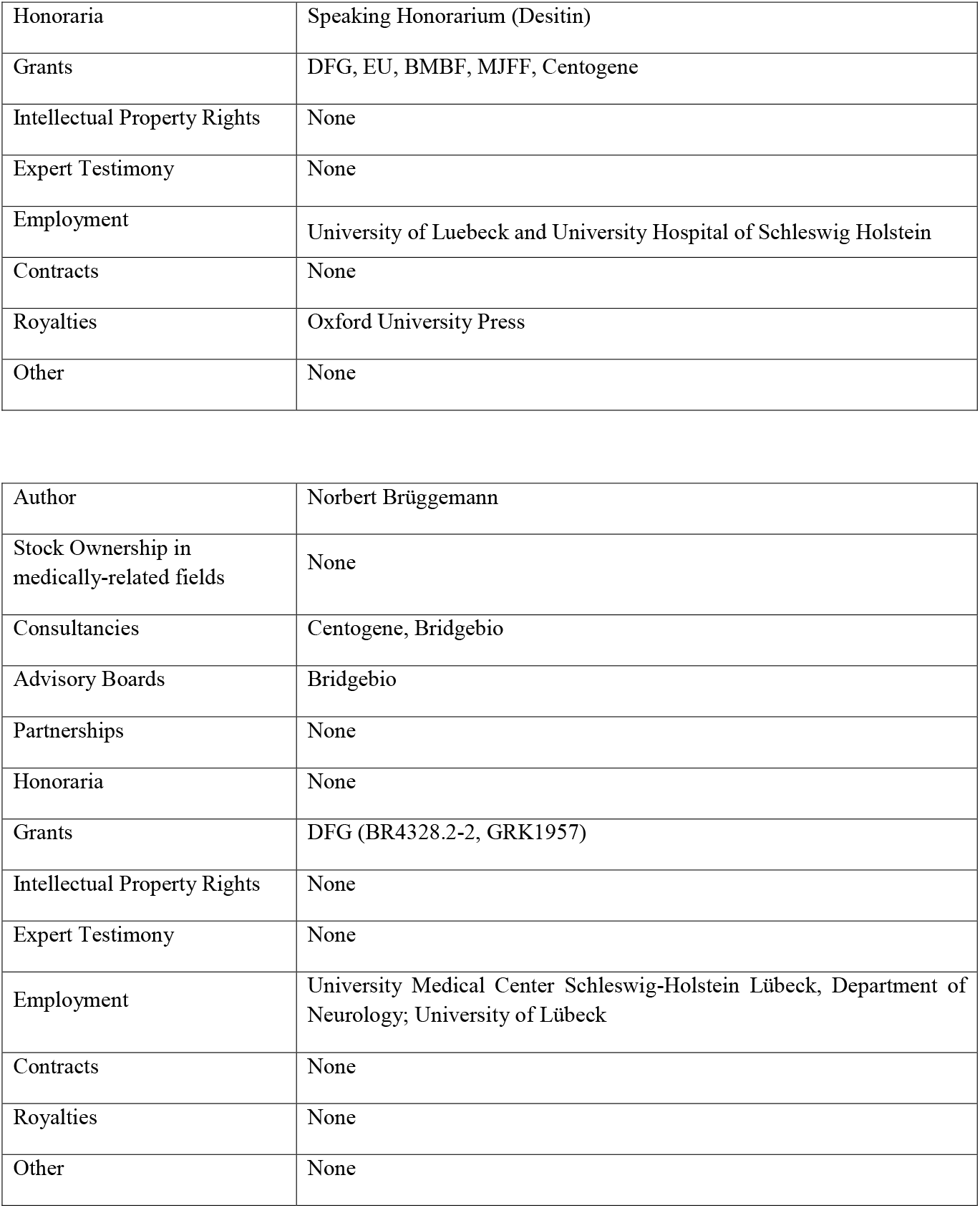

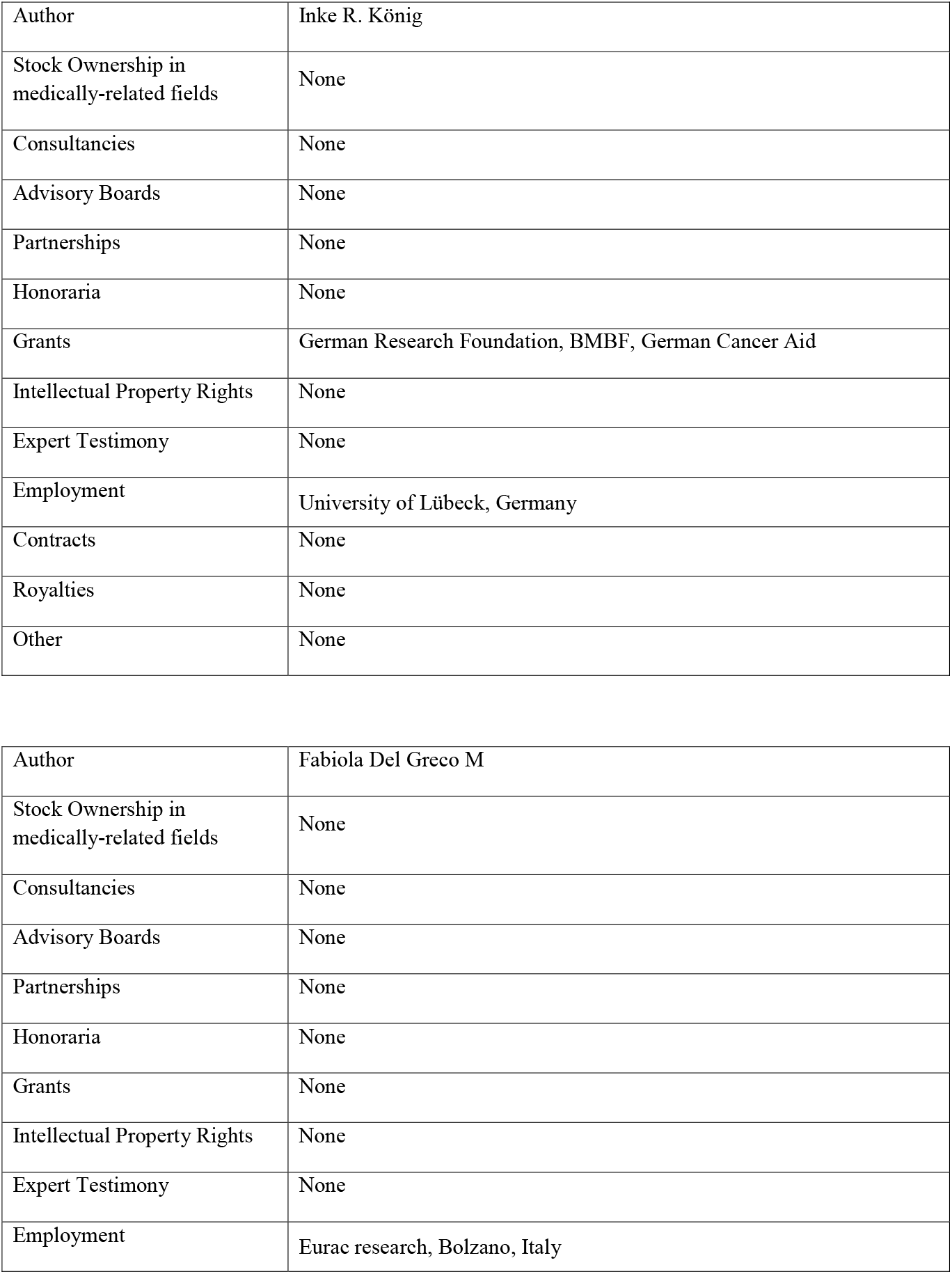

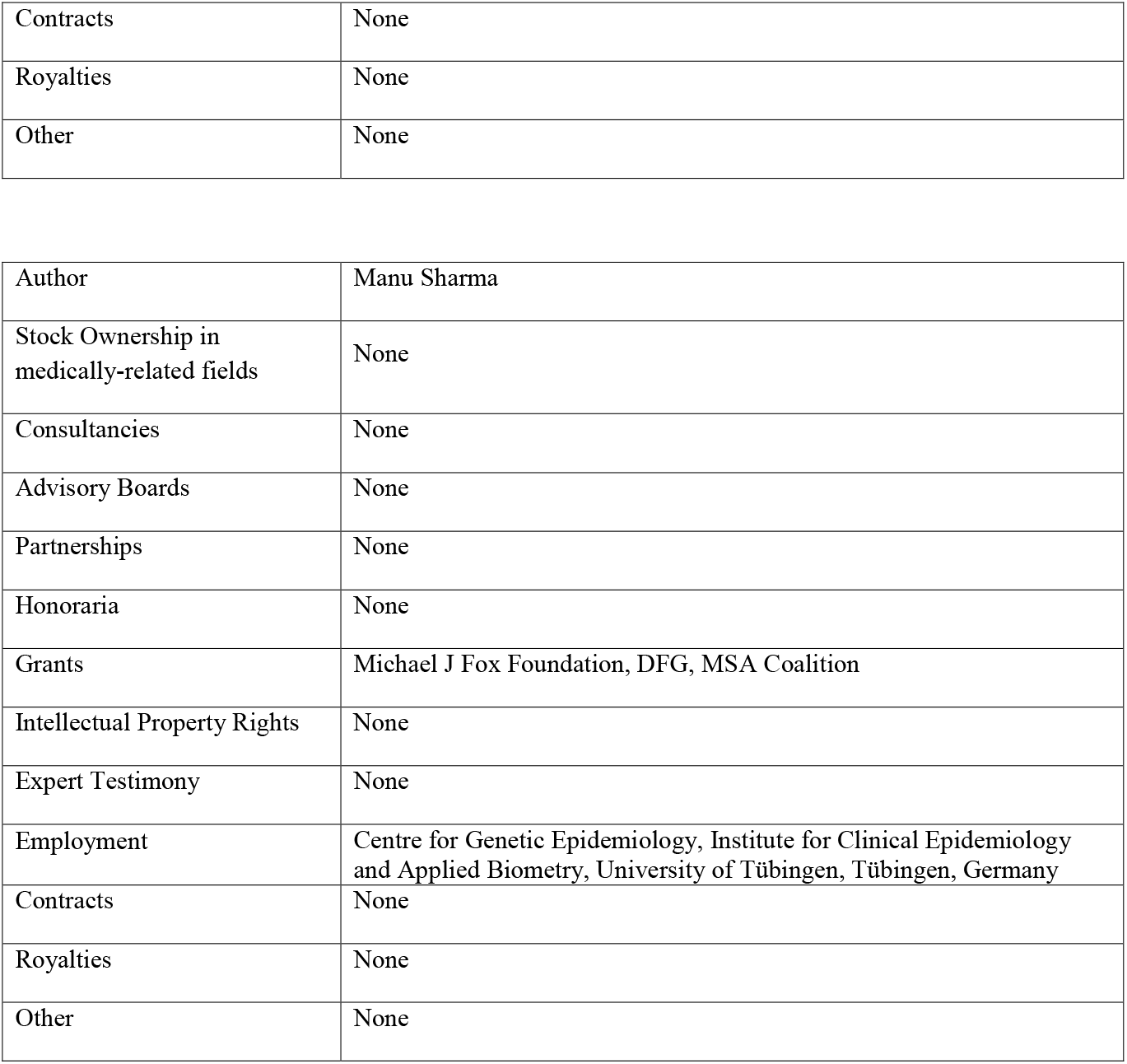

